# A novel method for robust estimation of ants’ walking speed and curvature on convoluted trajectories derived from their gait pattern

**DOI:** 10.1101/2021.08.08.455044

**Authors:** Jibeom Choi, Woojoo Kim, Woncheol Song, Sang-im Lee, Piotr Grzegorz Jablonski

## Abstract

Accurate measurements of travel distance and speed are crucial for the analysis of animal movements. Measuring the movements of ants entails measuring the change in locations registered at time intervals. This process involves dilemma of setting the proper time window: a short time window is vulnerable to spatial errors in observation, while a long time window leads to underestimation of the travel distance. To overcome these difficulties, we propose a novel algorithm that successively interpolates two consecutive points of ant’s trajectory for a given time window by embracing the alternating tripod gait of ants. We demonstrate that this algorithm is more reliable compared to the conventional method of travel distance estimation based on the sum of the consecutive straight-line displacements (SLD). After obtaining speed estimates for a range of sampling time windows, we applied a fitting method that can estimate the actual speed without prior knowledge of spatial error distribution. We compared results from several methods of speed and curvature extracted from the empirical data of ant trajectories. We encourage empirical scientists to utilize the proposed methods rather than the conventional SLD method of speed estimation as this process is a more reliable and subjective selection of the sampling time window can be avoided.

## 1. Introduction

In studies of movement ecology, the measurement of the distance that an animal traveled is an indispensable procedure to deduce the speed of the animal^1–3^. Despite their importance, there is a fundamental difficulty to accurately measure the moving animal’s distance and speed. The position of the animal is recorded discontinuously with certain intervals. For example, 30-fps video data provide the position of objects for every 1/30 sec, and the GPS position tracker updates its location information with a certain interval ranging from milliseconds to minutes.

Typically, researchers calculate the travel distance as a sum of the straight-line displacements (SLD; red dotted line in Fig 1a) along a path of a moving individual^4^. This leads to underestimation of the travel distance for low sampling frequencies (long sampling time window between consecutive measurements) even though there is no error in the measurement. The problem is more pronounced if the trail is more curved (shows higher tortuosity). A similar issue was considered by Benhamou (2004)^5^: low recording frequency of the individual’s positions on its walking path brings about a decrease in the measured path length, but shortening the (sampling) time window is vulnerable to the error, as explained below.

**Figure 1.**
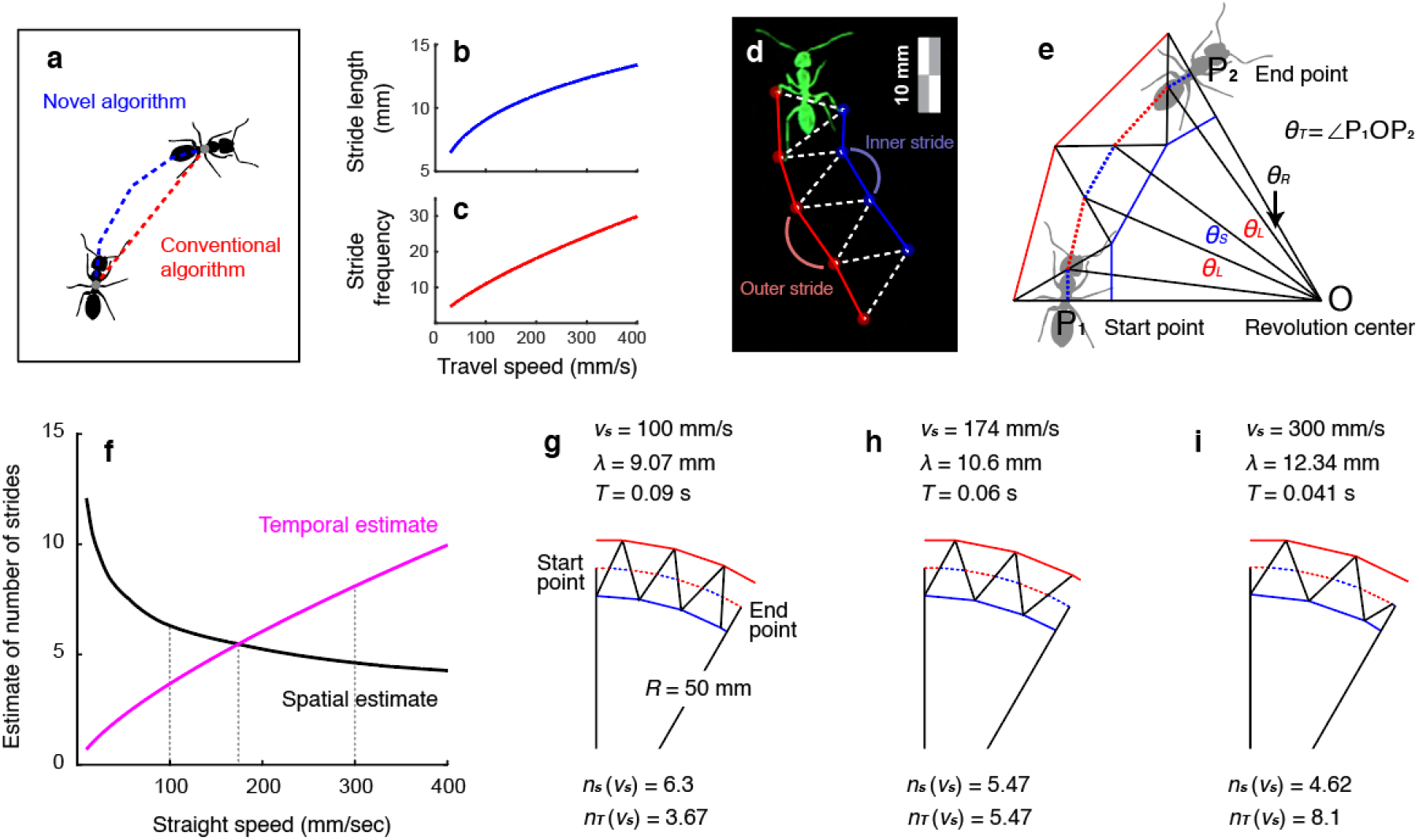
Basic assumptions and algorithms of novel speed estimation. **a**. Schematics of the difference of distance estimation between the conventional algorithm (sum over SLD) and the novel algorithm. As a novel algorithm considers the general pattern when making a turn, the distance estimated from the novel algorithm is always greater than that from the conventional one. **b-c**. The stride length should be nonlinearly proportional to speed (**b**, 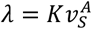) as well as stride frequency (**c**, 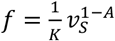). **d**. Analysis of ant gait from high-speed video by separating strides into the composition of triangles. The outer stride length (marked with a red line) is longer than the inner stride length (marked with a blue line). The color has been modified due to video processing of background elimination. See Supplementary Animation S1 for more details. **e**. Geometric analysis of stride number estimate based on angles generated by inner and outer strides. **f.** The estimate of the number of strides derived by spatial method (estimated from angles generated by inner and outer strides) and temporal method (estimated from window length divided by stride period) against straight speed given that radius of the circumscribed circle is 50 mm, the time window is 1/6 sec, and angle between start point and the end point is 30°. Specifically, the spatial method in this example assumes that the body center at the start point was in the middle of the outer stride. The spatial estimate (based on stride length) of the stride number is decreasing and the temporal estimate (based on stride period) of stride number is increasing as straight speed increases. The estimate of the straight speed is the speed where both stride estimates intersect. In this example, the estimate of the straight speed is 174 mm/sec. The spatial and temporal estimates when speed is 100 mm/sec, 174 mm/sec, and 300 mm/sec are represented in g, h, and i, respectively. **g**. The interpolation linking start point and the end point when straight speed is 100 mm/sec. The start point is on the left and the distance to the revolution center is 50 mm. The solid red and blue lines indicate strides of outer and inner strides, respectively. The dashed red and blue lines indicate the movements of the body center by outer and inner strides, respectively. In this case, stride length is 9.07 mm and stride period is 0.09 sec during which an ant performs an inner stride and an outer stride. Considering the angle between the start point and the end point around the revolution center (30° in this example), the spatial estimate of the number of strides is 6.3. Considering the length of the time window, the temporal estimate of the number of strides is 3.67. **h**. Same as (g) for straight speed of 174 mm/sec. In this case, the spatial and temporal estimates are identical, thus this straight speed is the estimated straight speed of the novel algorithm. **i**. Same as (g) for a straight speed of 300 mm/sec.

The problem of the shortened measured length would be ameliorated by estimation under short time windows, but this is vulnerable to the spatial error of the location measurement. Suppose that *δ* is the extent of error when measuring the location of the individual’s body center. If the actual length between *m* frames is *L*, the following holds:

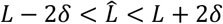

where 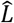 is the estimated length of *L*. Note that 2*δ*, not just *δ*, has to be added and subtracted as the measurements were performed at two points. Then the estimated speed is

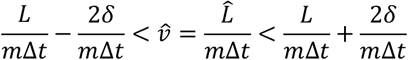

where Δ*t* is the time duration of a frame, and *m* is the frame difference in an integer unit. Written in another way,

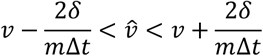

where *v* is *L* divided by *m*Δ*t* which would be close to the actual speed of an individual if *m* is small. As *m* becomes smaller, the proportion of the error 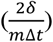 becomes larger, leading to inaccurate estimation. The short time window also produces a jagged estimated trajectory whose total length would be longer than the actual continuous one. Therefore, there is a tradeoff in the selection of time window length: the speed estimation using a longer time window is vulnerable to the effects of walking path tortuosity on estimates of speed, while that using a shorter time window is sensitive to errors.

With the advancement of automated multiple object tracking tools (e.g. AnTracks, idTracker.ai), researchers can obtain massive amounts of ant position data from video recordings^6–10^. Even if there is no error in measurement, however, linking the body centers between frames would lead to a shortening of the measured travel distance compared to the actual travel distance on a curved trajectory. Although shortening of the sampling time window may help with this issue, it eventually leads to more pronounced errors in estimation due to spatial noise in the estimation of the location of the two points. The dilemma of time window selection was discussed by Wang & Song (2016)^11^ explaining that a short time window leads to unstable fluctuation of data and a long time window disturbs the analysis of the detailed configuration of the trail. As the effect of the error can be attenuated by longer window length, and underestimation in the measurement can be ameliorated if trails are straight, many ant studies measure speed only on the straight fraction of trajectory without abrupt stop or acceleration^11–14^. There have been numerous attempts to resolve this issue by building specific models and using specific assumptions. These models require specific information about the trajectories such as error distribution^15,16^ or require specific stochastic assumptions^17,18^.

In order to reliably estimate the speed and curvature of ant movements from observations, we developed an algorithm that uses the alternating tripod gait pattern of ants^14,19,20^ to interpolate two consecutive points on a trajectory. The algorithm takes into account the positive correlations between ant’s speed and stride frequency as well as between ant’s speed and stride length which have been previously clarified by empirical studies^14,21,22^. These traits that are specific to ants allowed us to develop methods of ant speed estimates which are proven to be relatively accurate when there are errors in observation. We also introduce the method that infers the speed and optimal sampling window length by looking into the relationship between observed speed and sampling window length. This approach might be applicable to other hexapod species (or robots) if they follow similar pattern of alternating gait and strategy of turns.

## 2. Materials and Methods

### 2.1. Algorithm Explanation

#### 2.1.1. Stride frequency and length

As stated above, it has been found that there is a positive correlation between travel speed and stride frequency as well as between travel speed and stride length^13,14,21,22^. While linear regressions are often assumed in the literature^13,14,21^, assuming linear regression between travel speed and stride frequency and that between travel speed and stride length at the same time is logically contradictory. Let us assume that frequency and stride length are expressed as follows:

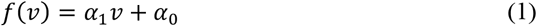

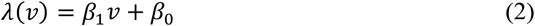

where *f λ, v* are the stride frequency, stride length, and speed of ant following a straight trajectory, respectively. It is evident that *v* = *f*(*v*)*λ*(*v*) should hold regardless of *v*. This is because speed is equivalent to the travel distance during a time period divided by the length of the time period, and the inverse of the time period is frequency. (The same rule is found in optics that the speed of the wave is the product of wavelength (analogous to the stride length) and frequency.) Consequently, assuming equations (1) and (2) are true, we expect that the following should hold as well:

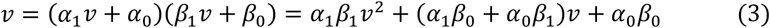

As *α*_1_ and *β*_1_ are non-zero (as the positive correlation is assumed), the equation is quadratic. Hence, there are at most 2 solutions of *v* that satisfy the equation. It is, therefore, impossible for the equation (3) to hold for every *v* indicating that the relationships postulated in equations (1) and (2) are not realistic.

To resolve this issue, we suggest using other formulas for stride frequency (*f*(*v*)) and stride length (*λ*(*v*)) against speed, which are

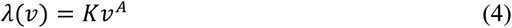

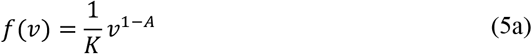

so that *f*(*v*)*λ*(*v*) = *v* holds (Fig. 1b, c).

As the frequency (*f*(*v*)) is defined as the inverse of the time period, equation (5a) can be represented as equation 5b below:

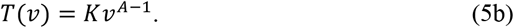

As a period in physics is defined as the span of time between successive occurrences of the identical state in a cyclic movement, we defined stride period (*T*) of gait in this paper as the span of time when the ant performs a full single gait including left and right strides. On a curved trajectory, an ant performs outer and shortened inner stride during a stride period (Supplementary Video S1).

#### 2.1.2. Model Structure

According to Zollikofer et al. (1994)^22^, ants perform alternating tripod gait along straight and curved paths with conserved schematics of tripod configuration. When ants turn, they shorten the length of the inner stride (i.e. the stride that lies in the inner side with respect to the revolution center; Supplementary Video S1, Fig. 1d). Inspired by these empirical findings, we built a model that simulates ants performing turns by asymmetrical alternating tripod gait. Since the length of the outer stride in a turn is the same as the stride length when ants walk straight, we denoted the length of the outer stride as *λ*. The length of the inner stride is shortened by some extent *kλ* (0 < *k* ≤ 1). This shortening of the inner stride length is observed in our video analysis as well (Supplementary Video S1).

In our algorithm, we represented the ant’s alternating tripod gait as a series of triangles. The triangles are defined by the contact points of the ground (Supplementary Video S1) and the tarsi of two middle legs (L2 and R2). The example of those triangles is described in the still image (Fig. 1d, e; Appendix Fig. A1–A3). For sake of simplicity, triangles were assumed to be isosceles. There are two types of triangles: those with the base of length *λ* (henceforth, triangle *Ω_L_* whose base is longer stride (red line in Fig. 1d, e)) located on the outer stride of the turn, and those with the base of length *kλ* (henceforth, triangle *Ω_s_* whose base is shorter stride (blue line in Fig. 1d, e)) located on the inner stride of the turn. The height of *Ω_s_* was assumed to be equivalent to the span of the ants’ legs during walking. Using this framework, we described the movement of the body center as the combination of two distinct linear parts: the movement gained from the outer stride and from the inner stride. As *Ω_L_* and *Ω_s_* are isosceles, the travel distance of the body center is *λ*/2 and *kλ*/2 for outer and inner stride respectively. Three consecutive points of observation (P_1_, P_2_, P_3_) separated by certain time window duration (*Δt*) were used to deduce the curvature radius (*R*). The speed of the body center is the average of values calculated for four specific moments (henceforth referred to as body center condition). Each value is derived from the moment when body center is (1) at the middle of the outer stride; (2) at the middle of the inner stride; (3) at the beginning of the outer stride (at the end of the inner stride); (4) at the beginning of the inner stride (at the end of the outer stride). If the body center moves at a constant speed during each of the strides and the durations of inner and outer strides are equal, the probability that the body center is measured in each of the four aforementioned conditions would be the same. In the first round of the calculation, *P*_1_ is the start point and assumed to satisfy one of the four aforementioned conditions. The endpoint is P_2_ and the stage of the gait in the point is not specified. In the following round of the calculation, P_2_ is the start point and assumed to satisfy one of the four conditions. The endpoint is P_3_ and the stage of the gait is not specified. The speed of trajectory from P_1_ to P_3_ is the average speed calculated from these two rounds of calculations (distance estimation from P_1_ to P_2_, and that from P_2_ to P_3_). See Appendix for the details.

#### 2.1.3. Estimation of straight speed (v_s_) from temporal and spatial estimation of stride numbers

We propose a method of estimation of the straight speed (*v_S_*) based on the estimates of the number of strides (*n_S_*) performed in a time window (*Δt*). Straight speed (*v_S_*) is the hypothetical speed at which the ant would have walked without shortening of the inner stride (*kλ*) relative to the outer stride (*λ*). As ants shorten the inner stride length during the turn, the actual speed of body center (*v_c_*) is always slower than straight speed. First, the straight speed (*v_S_*) needs to be determined. From the previous assumptions on stride length (*λ*), stride frequency (*f*), and stride period (*T*), it holds that 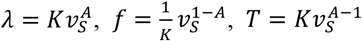. Suppose that two consecutive positions of ants along the curved track with revolution radius of *R* are given by their body center positions (P_1_, P_2_; Start point and end point in Fig 1g, respectively) recorded at the temporal difference of *Δt* (length of sampling time window) from each other. *R* is determined by the circumscribing circle generated by P_1_, P_2_, and P_3_. Suppose that straight speed (*v_S_*) is fixed, and thus so is outer stride length (*λ*). Then, it is possible to deduce the angles generated by the movement of the body center during the outer and inner stride (*θ_L_* and *θ_S_*, respectively; Fig. 1e)) with respect to the revolution center (O, the center of the osculating circle). The travel distance of the body center during the stride is the half of the stride length. Additionally, the longer the revolution radius is, the smaller the *θ_L_* and *θ_S_* are, given that stride length is fixed. The number of the outer and inner strides should satisfy the condition that *n_i_θ_L_* + *n_o_θ_S_* = *θ_T_* = ∠P_1_OP_2_ where *n_i_, n_o_* are the number of the inner and outer strides, respectively, and *θ_T_* is the total angle around the revolution center generated by the movement of an ant during a time window. The specific geometric procedure of estimating *n_i_, n_o_* are presented in the Appendix. The number of the total strides (*n_S_*) is calculated as the sum of inner and outer strides (*n_S_* = *n_i_* + *n_o_*). This number of strides calculated from a given angle *θ_T_* can be expressed as a function of straight speed (*v_S_*). As stated previously, straight speed is assumed to determine the outer stride length (*λ*), frequency (*f*), and period (*T*) of the gait. The function *n_S_*(*v_S_*) then determines this number of strides against straight speed (*v_S_*), and this number is the spatial estimate of the stride number in Fig 1f, black line).

On the other hand, the total number of strides can be also calculated from temporal aspects of walking gait. Within the temporal window length of *Δt*, the number of the strides can be calculated from the stride period. As two strides (inner and outer) are performed during a single period, the total number of the strides during the window length is *n_T_* = 2*Δt/T* and period is determined by straight speed as already mentioned 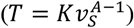. When Δt is constant, *n_T_* is solely determined by straight speed. Let this number of strides calculated from the stride period be expressed as a temporal number (temporal estimate; Fig. 1f, pink line), which is a function of walking speed (*n_T_*(*v_s_*)).

Given that one of the four aforementioned body center conditions at the recording is fulfilled, it is possible to derive the single specific value of 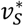 such that 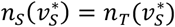 holds. This is validated by the following properties of *n_S_* and *n_T_*: (1) *n_S_* is monotonically decreasing and *n_T_* is monotonically increasing against *v_S_* (Fig. 1f); (2) *n_S_*(*v_S_*) – *n_T_*(*v_S_*) > 0 for very small *v_S_* (Fig. 1f and an example in Fig 1g) and *n_S_*(*v_S_*) – *n_T_*(*v_S_*) < 0 for very large *v_S_* (Fig. 1d and an example in Fig 1g). From properties (1) and (2), one and only one specific 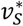 is determined by finding a situation when *n_s_* = *n_T_* (Fig. 1f and an example in Fig. 1h). Hence, 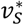 is the estimate of the straight speed derived from simultaneously considering spatial and temporal evaluations of the number of strides. Fig. 1f-i illustrate process of identifying 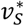. If 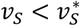 (Fig. 1g, *v_S_* = 100 mm/sec), then *n_S_*(*v_S_*) is greater than *n_T_*(*v_S_*), and *vice versa* if 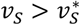 holds (Fig. 1i, *v_S_* = 300 mm/sec). The two estimates of the stride numbers (the spatial estimate and the temporal estimate) are identical when the straight speed *v_S_* =174 mm/sec (Fig. 1h), which is the estimate of the straight speed for Fig. 1f.

After obtaining 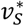, the *λ* and *kλ* are calculated in order to estimate the actual speed of the body center in a curved trajectory (*v_C_*). Detailed mathematical methods are presented in the **Appendix**. Considering the shortening of the inner stride, the actual body speed for the condition of Fig. 1f, h (where 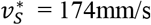) is *v_C_* =157 mm/sec.

### 2.2. Testing the new method with simulated walking paths

#### 2.2.1. Generation of simulated trajectories

To test the robustness of the novel algorithm of speed estimation presented in this study, we ran five types of simulations that mimic the movement of the ants. In all of the simulations, we assumed that there are errors when estimating the position of the ants’ body center. The width and length of ants’ torso (body without legs) are expressed as *W_B_* and *L_B_*, respectively in this model. Suppose that the body axis of an ant is located along the *y*-axis and its center is located at the origin in the two-dimensional Cartesian coordinate system. The observed position of the ant center is expressed as follows:

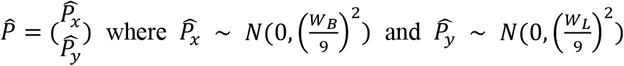

According to the traits of normal Gaussian distribution, 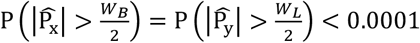.

In *Simulation 1*, an ant in a simulation moves in a linear trajectory. A random point in a trajectory was selected and two more points after *Δt* (observation window length) and 2*Δt* were additionally collected with the observation errors mentioned above. Denote by those three points as 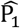 (randomly chosen point), 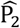 (the point after *Δt* compared to 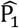) and 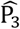 (the point after 2*Δt* compared to 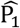), respectively.

The conventional method to estimate the speed is as follows:

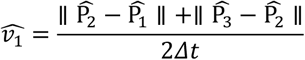

where ║·║ represents Euclidean distance between two points.

The novel method to estimate the speed is as follows:

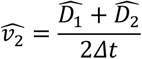

where 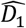 and 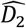 are the distance that an ant moved estimated by the proposed algorithm considering the actual gait pattern of ants (the details of the algorithm are explained in *2.1. Algorithm Explanation*). We also measured the radius of circumscribed circle generated by the three points.

In *Simulation 2* and *3*, ants move along the curved path (circular in Fig. 2a or sinusoidal-like in Fig. 2b). In each stride, the body center moves linearly, but the direction is modified at the end of the stride. We set the maximal distance from the revolution center to the body center to be *R_m_* (Fig. 2a, b; *R_m_* = 50 mm for Simulation *2.1* and *3.1*; *R_m_* = 30 mm for Simulation *2.2* and *3.2*). At the end of each advance by the inner or outer stride, the body center meets the circle with a radius *R_m_*. (The definitions of inner and outer strides are presented in *2.1. Algorithm Explanation*). This circle defining the outermost position of the body center is fixed for *Simulation 2* so that the ant revolves around a certain point. In *Simulation 3*, on the other hand, the orientation of the revolution (clockwise or counterclockwise) alternates at the end of each semicircle. The ant advances with periodic waves composed of semicircles. The direction of movement was considered when deriving errors of measurement on assumption that the body axis is always parallel to the movement direction. As in *Simulation 1*, we recorded the speed obtained by the conventional and novel algorithm and radius of circumscribed circle generated by the three points. This observed radius was compared with the actual radius of the trajectories to evaluate the reliability of the measurement (*2.2.3. Measuring the reliability of traditional and novel methods applied to simulated paths*). The actual radius of the trajectory was calculated as the mean distance from the polygon center.

**Figure 2.**
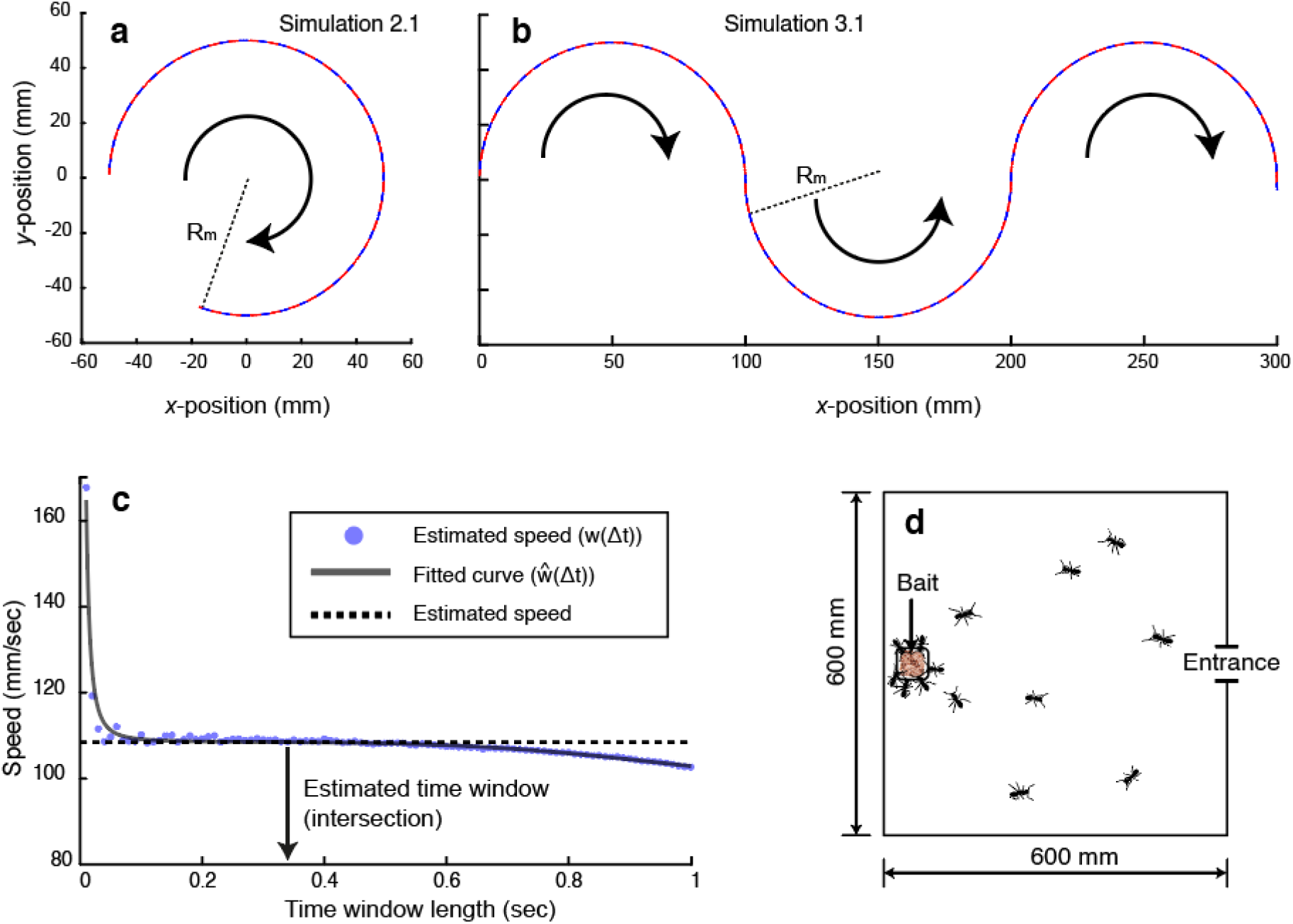
The structure of simulations and empirical experiments. **a**. The trajectory of *Simulation 2.1*. Ants walk around a circular trajectory, and the body center is propelled by long stride (outer stride, red) and short stride (inner stride, blue). **b**. The trajectory *Simulation 3.1*. Ants walk along the sinusoidal-like path composed of semicircles. **c**. The estimation of the speed through the fitting method. The fitted equation includes the term generated by errors and sinuosity. The data points are obtained novel algorithm in *Simulation 3.1*. **d**. We placed 600 mm × 600 mm experimental arena with bait (grinded tuna) on three different colonies (AG, MS, MG) and recorded movements of ants.

The expected radius by the outer stride is obtained as follows:

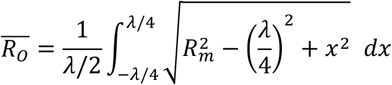

where *λ* is the stride length of the outer stride. It was assumed in the model that the body center moves the distance of *λ*/2 by the stride.

The expected radius by inner stride is obtained as follows:

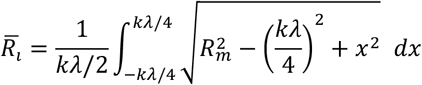

where *kλ* (0 < *k* ≤ 1) is the stride length of the inner stride.

As ants in a simulation move at a constant speed in each stride with the same temporal period, the expected radius among the movement is 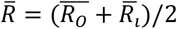. We compared this value with the estimated radius calculated from the circumscribed circle by the index of geometric curvature (1/*R*) which was also presented in Zollikofer (1994).

#### 2.2.2. Fitting of walking path data for speed estimation

We applied the procedures explained in *2.1.2 and 2.1.3* to determine the walking speed and geometric curvature for different sampling windows (*Δt*; blue dots in Fig. 2c represent such estimates). When the sampling time window is short, the expected value of the estimated path length would exceed the real path length due to the effect of the observation error and corresponding crinkles of the observed path. For example, suppose that there are observation errors in a linear trajectory. The estimated path is wiggled and crinkled due to errors so that the estimated distance is generally greater than the actual distance. This effect of the error would be mitigated if the window length is extended, which would elicit another kind of problem explained below.

If ants perform random walking, then the ratio of the distance between two points to the actual travel distance between these points tends to zero as the time window gets longer^5^. If ants perform directed walking with conserved tortuosity, then the ratio would converge to some value as the time window gets longer. As a generalized approximation, the distance between two points can be expressed as ((1 – *B*)*Ψ*(*Δt*) + *B*)*vΔt* where 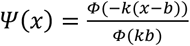 and *Φ*(*z*) is the cumulative distribution function of the standard normal distribution. By this structure, (1 – *B*)*Ψ*(*Δt*) + *B* is 1 when *Δt* = 0 and (1 – *B*)*Ψ*(*Δt*) + *B* tends to *B* as *Δt* gets larger for positive *k. B* = 0 for a perfectly random walk as measured distance between two points compared to actual length becomes 0 as *Δt* gets larger^5^. *B* = 1 for a perfectly straight walk as it is possible to measure the almost accurate length of the straight trajectory if the time window is long. Generally, 0 ≤ *B* ≤ 1 should hold based on the tortuosity of the trajectory. For example, *B* should be 1 and 0 for *Simulation 1* and *2*, respectively. *B* should be 2/*π* for *Simulation 3* because the ratio between the measured distance between two points and actual distance becomes ratio between diameter (2*R*) and circumference of the corresponding semicircle (*πR*) if the time window is infinitely long.

We made a fitting equation based on these two effects of error and tortuosity. Applying the Pythagorean Theorem, the measured distance between two points can be approximated as 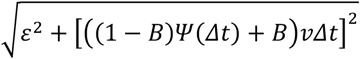 where *ε* is the effect of increased measured distance due to measurement error. The estimated speed by time window (*w*(*Δt*)) is the measured distance (by SLD or novel algorithm) between two points divided by the time window length (*Δt*) which is a function against the time window. If the novel algorithm of the speed estimation was used, then *w*(*Δt*) = *v_C_* holds.

The examples of *w*(*Δt*) in simulation and empirical data are represented as scatter plots in Fig. 2c and Fig. 4c3-e3, respectively. Based on prior arguments, function *w*(*Δt*) can be fitted into the following theoretical equation (*ŵ*(*Δt*)).

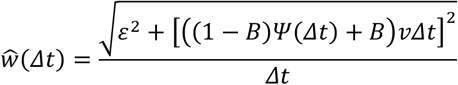

**Figure 3.**
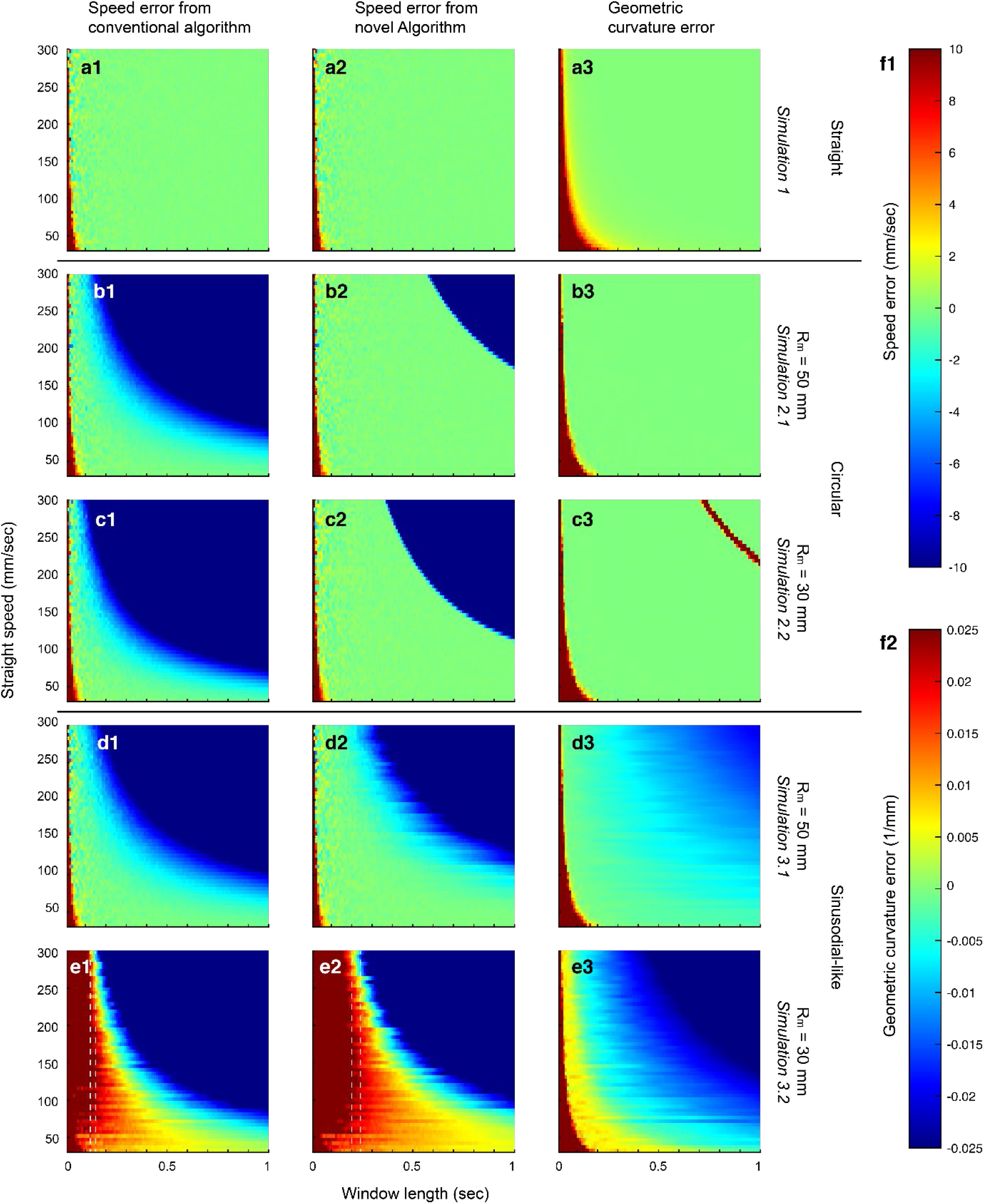
The heatmaps representing errors in speed (the first and the second column) and errors in geometric curvature. **a1-a3**. The error heatmaps for *Simulation 1* where the actual trajectory is linear. **a1** represents the error heatmap obtained by the conventional algorithm, and a2 represents the error heatmap by the novel algorithm. **a3** is an error heatmap of geometric curvature. **b1-b3**. Same as **a1-a3** for *Simulation 2.1* where the actual trajectory is circular with a maximum radius of 50 mm. **c1-c3**. Same as **a1-a3** for *Simulation 3.1* where the actual trajectory is composed of alternating semicircles with a maximum radius of 50 mm. **d1-d3**. Same as **b1-b3** for *Simulation 2.2* with a maximum radius of 30 mm. **e1-e3**. Same as **c1-c3** for *Simulation 3.2* with a maximum radius of 30 mm. Vertical dashed lines indicate the reliable band of time window length within which the error value is generally less than 10 mm/sec. **f1**. Error legend of the heatmap for speed error. **f2**. Error legend of the heatmap for geometric curvature error. As explained in *2.2.3*., there is no difference in the estimation of geometric curvature for conventional and novel methods.

**Figure 4.**
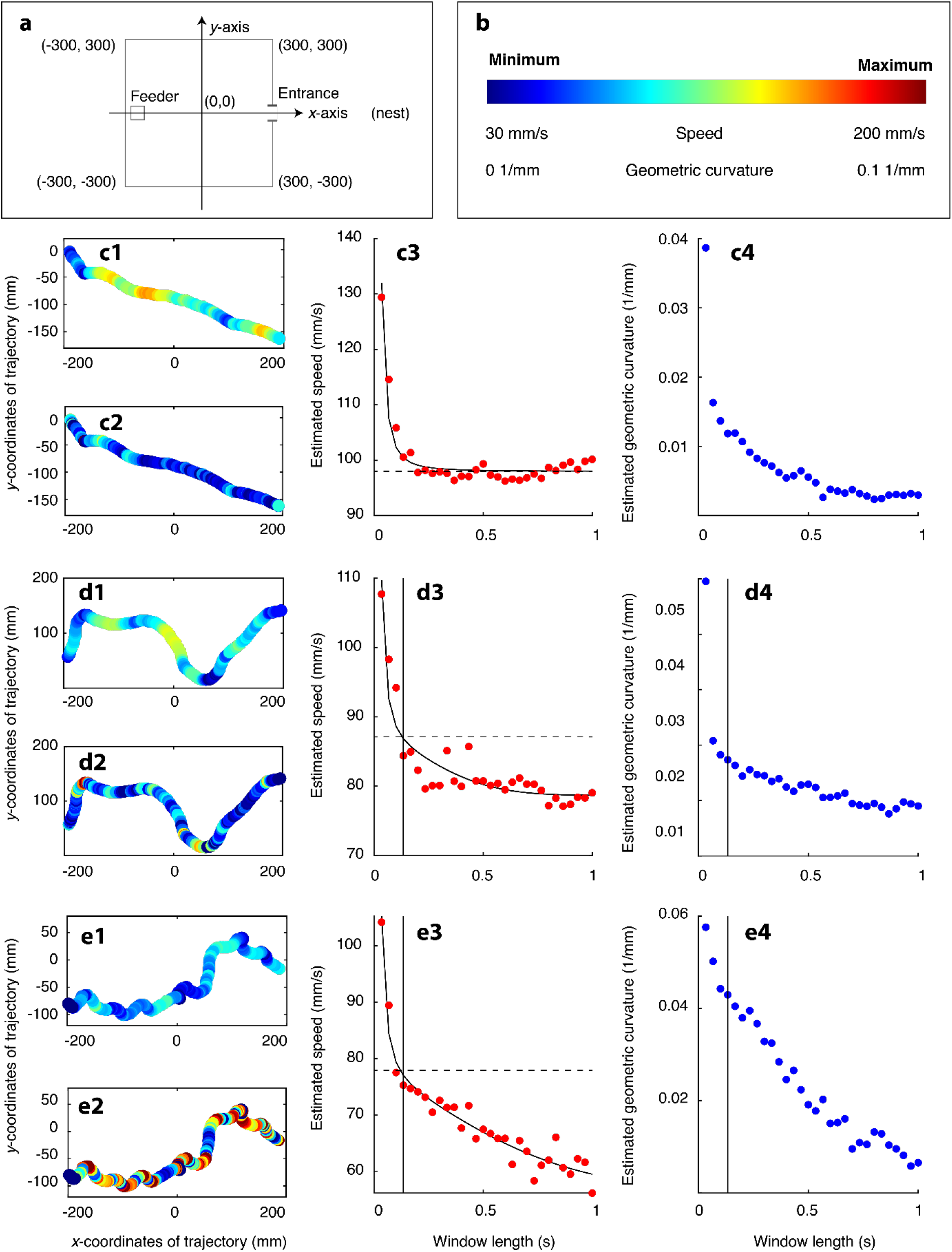
Examples of trajectories gathered from empirical experiments and corresponding analysis results. **a**. Coordinate system of the arena. **b**. Legend of speed and geometric curvature for trajectory description for **c1, c2-e1, e2**. **c1**, Trajectory colored by the speed estimated with 5 frame window length. **c2**. Trajectory colored by the geometric curvature with 5 frame window length. **c3**. The speed estimated with different time window lengths from an example trajectory (**c1**). The best-fit equation marked with the solid curve describes the relationship between the sampling time window length and the empirically determined speed. The theoretically estimated speed is marked with the horizontal dashed line. The point where the fitted equation and the horizontal speed intersects provides information about the optimal sampling window length (in this case 1 sec). **c4**. The geometric curvature estimated with different time window lengths from an example trajectory **c1**. The proper value of geometric curvature was determined by the value estimated from window length in **c3**. **d1-d4** Same as **c1-c4** for another trajectory. In this case, the time window where fitted curve and estimated velocity meet is represented with vertical solid line (0.1 s). **e1-e4**. Same as **c1-c4** for another trajectory.

By fitting this equation into the observed speed with different time windows, one can deduce estimated values of *ε*^2^, *B*, and *v*. The genuine value of speed would be this *v* estimated from nonlinear regression.

#### 2.2.3. Measuring the reliability of traditional and novel methods applied to simulated paths

The observed speed against time window (*w*(*Δt*)) for each trajectory can be obtained in two ways: one with the conventional method (consecutive straight-line displacements (SLD); Euclidean distance divided by time) and another with the novel algorithm (Distance attained by the proposed algorithm divided by time). We obtained w(dt) for both methods. As novel method was performed by procedures explained in 2.1.3 (Fig. 1) for a series of time windows, this resulted in specific speed values for specific *Δt* (Fig. 2c and Fig. 4c3-e3). Then we estimated speed by the aforementioned fitting methods (the dashed horizontal line in Fig. 2c2c and Fig. 4c3-e3). An alternative way of deducing speed is to set the specific time window. Regardless of the shape of *w*(*Δt*), one can estimate the speed by, for example, assuming that a reliable time window is 0.1 sec. Then, the estimated speed is *w*(0.1).

To sum up, there are two methods of obtaining the relationship between the speed and time window length, *w*(*Δt*), and there are other two methods of estimating speed out of this relationship (*w*(*Δt*)). Observed speed can be obtained by simply linking the body center over a distance between two consecutive observation points separated by a certain time window (SLD), or it can be obtained by applying a novel algorithm proposed in this study. Let these two methods be denoted by conventional and novel algorithms, respectively. After obtaining *w*(*Δt*), one can estimate speed by setting the fixed time window, or it can be estimated by nonlinear regression. Let these two methods be denoted by point estimation and fitting estimation, respectively. Both methods of speed estimation can be applied to the two results of distance estimation (conventional SLD and the novel algorithm considering the gait pattern of the ants).

Similarly, geometric curvature defined by 1/*R* where *R* is the radius of the circumcircle generated by the three points (triplet) also varies with the time window and this can be estimated by two methods. One method is to get the geometric curvature with a fixed time window. Another method is to retrieve the optimal time window (*Δt*\*) that satisfies 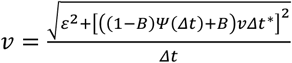 for estimated *ε*^2^, *B*, and *v*. The fitted observed speed measured from the optimal time window *ŵ*(*Δt*\*) equals the genuine speed value. This is the point where the fitted function and horizontal line indicating genuine speed intersect in Fig. 2c and Fig. 4c3-e3. Then, the estimated geometric curvature is the one that corresponds to the optimal time window (vertical lines in Fig. 4d3, d4, e3, e4).

There is no difference in point estimation of geometric curvature for observed speed of conventional and novel methods because there is no difference in observed geometric curvature by time window for the two methods.

For each of the five simulation results (straight path, circular paths with maximal radius of 50 mm and 30 mm, sinusoidal-like paths with maximal radius of 50 mm and 30 mm), we estimated speed and geometric curvature by eight methods (FitC, FitN, PtC_0.1_, PtN_0.1_, PtC_0.3_, PtN_0.3_, PtC_0.5_, PtN_0.5_ (The subscripts of point estimation method (Pt) refers to the fixed time window length); Table 1). We compared these estimates from eight methods to the ‘real’ values of simulated paths. In each simulation, the deviation of the measurement for each method (Θ_*i,j*_ for *i*-th simulation and *j*-th method, 1 ≤ *i* ≤ 5, 1 ≤ *j* ≤ 4) is defined as

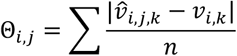

where 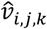 stands for estimated speed using *j*-th method for *k*-th speed value in *i*-th simulation and *v_i,k_* is the corresponding real speed. We confined the straight speed of the simulation to the range between *v_i,k_* = 30 mm/sec for *k* = 1 to *v_i,k_* = 200 mm/sec for *k* = 55 as the measured speed of empirical data rarely exceeded 200 mm/s and speed slower than 30 mm/sec was considered to be stationary. The fixed time window length was set to be 0.1 sec as this window length exhibited higher accuracy based on *Simulation 2.1, 3.1*, and *2.2*.

**Table 1.**
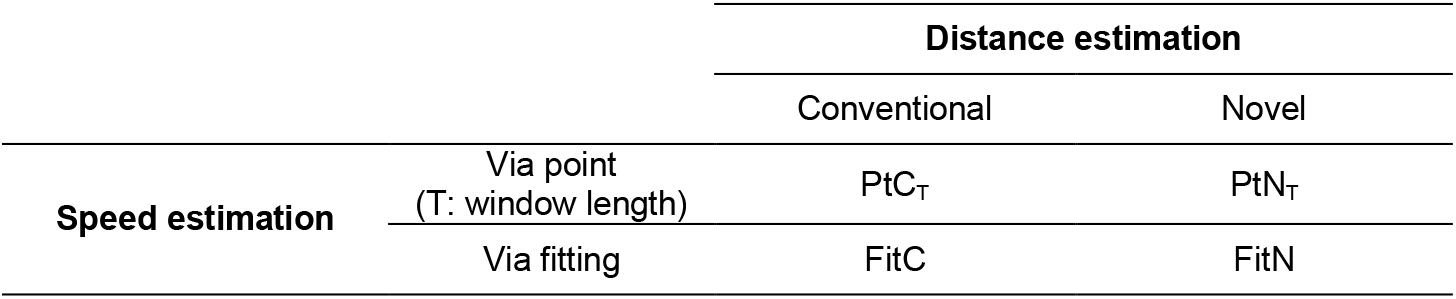
The classification of estimation methods according to distance estimation and speed estimation approach. Speed observation has two levels of conventional algorithm and novel algorithm. The conventional algorithm estimates distance by the sum over the straight-line displacements (SLD) which is frequently used by common ant research. The novel algorithm utilizes the actual gait pattern of the ants. When estimating speed, the window length for the point method is fixed for 0.1 sec while that of the fitting method is determined by schematics of the observed speed against the time window.

To standardize the scale of errors for each simulation, the index of standardized deviation (*θ_i,j_*) was defined as

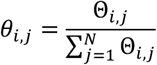

After obtaining the index for each simulation and method, we used the mean of these indices for five simulations as the measure of reliability. We recorded the standardized deviation of the speed and geometric curvature estimation for each simulation. For each of the empirically recorded trajectories, we estimated their speed and geometric curvature using eight methods (see Table 2 and 3 for results of each estimation) and compared them.

**Table 2.** The mean standardized deviation (used as an index of the ‘degree of error’ in estimation of real values) for each algorithm in each simulation for speed estimations. Explanations of all methods are presented in Table 1.

|  | Speed |  |  |  |  |  |  |  |
| --- | --- | --- | --- | --- | --- | --- | --- | --- |
|  | FitC | FitN | PtC<sub>0.1</sub> | PtN<sub>0.1</sub> | PtC<sub>0.3</sub> | PtN<sub>0.3</sub> | PtC<sub>0.5</sub> | PtN<sub>0.5</sub> |
| **S1** | 0.122 | 0.116 | 0.250 | 0.250 | 0.089 | 0.088 | 0.042 | 0.042 |
| **S2.1** | 0.024 | 0.013 | 0.034 | 0.014 | 0.248 | 0.004 | 0.661 | 0.003 |
| **S3.1** | 0.025 | 0.019 | 0.031 | 0.012 | 0.190 | 0.049 | 0.448 | 0.226 |
| **S2.2** | 0.023 | 0.011 | 0.025 | 0.005 | 0.201 | 0.002 | 0.502 | 0.231 |
| **S3.2** | 0.143 | 0.153 | 0.088 | 0.125 | 0.088 | 0.050 | 0.199 | 0.155 |
| **Mean** | 0.067 | 0.062 | 0.086 | 0.081 | 0.163 | 0.039 | 0.370 | 0.131 |

**Table 3.** The mean standardized deviation (used as an index of the ‘degree of error’ in estimation of real values) for each algorithm in each simulation for geometric estimations. Explanations of all methods are presented in Table 1. The estimates from PtC_T_ and PtN_T_ are the same so they are represented as Pt_T_.

|  | Geometric curvature |  |  |  |  |
| --- | --- | --- | --- | --- | --- |
|  | FitC | FitN | Pt<sub>0.1</sub> | Pt<sub>0.3</sub> | Pt<sub>0.5</sub> |
| **S1** | 0.009 | 0.009 | 0.890 | 0.067 | 0.025 |
| **S2.1** | 0.069 | 0.040 | 0.885 | 0.004 | 0.003 |
| **S3.1** | 0.087 | 0.145 | 0.276 | 0.186 | 0.307 |
| **S2.2** | 0.123 | 0.092 | 0.774 | 0.008 | 0.003 |
| **S3.2** | 0.152 | 0.136 | 0.202 | 0.192 | 0.318 |
| **Mean** | 0.088 | 0.084 | 0.605 | 0.091 | 0.131 |

### 2.3. Applying the novel methods to empirical data on ant trajectories

In order to provide an example of application of the novel methods to empirical data, we used data from field experiments (Fig. 2d). In three colonies of the Japanese carpenter ant (*Camponotus japonicus*) located at Seoul National University Gwanak campus, we placed 600 mm × 600 mm experimental arena on which ground tuna was piled up. These three colonies were the same colonies that were studied by Choi et al. (2020)^10^. We grabbed ants near the colony and placed them near the bait to generate trajectories from the bait to the nest. We filmed the formation of the trajectories for 100 minutes. 13 experiments were performed for each colony so that the total length of the video was 3,900 minutes (= 13 × 100 × 3 = (repeated experiments) × (duration of each experiment) × (the number of the colonies)). After video processing for the sake of tracking fidelity (Adobe Premiere Pro), we analyzed the video and extracted temporal and spatial coordinates of each trajectories using AnTracks (https://sites.google.com/view/antracks).

## 3. Results

### 3.1. Comparison of the conventional and novel algorithm in simulations

In order to visually represent the reliability of the two methods of speed estimation, we represented the difference between estimated speed 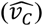 and the actual speed (*v_c_*, this value is known as we simulated the movements). We used a heat-map method (Fig. 3 a1-e1, a2-e2) to visualize the value of the difference between speeds from the two methods 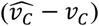 presented as the ‘speed error’ in Fig. 3. In all of the simulations, the actual speed is overestimated if straight speed and time window duration are low. On the other hand, it is underestimated if straight speed and time window length are high (Fig. 3b1-e1, b2-e2), except for the simulation of the ideally straight walk (Fig. 3a1, a2). For *Simulations 2.1* and *2.2*, where the ant trajectory is circular, the degree of error is similar for both algorithms if straight speed and window length are low.

When the measured distance between two observation points is long, (i.e., when speed and window length are high) the measured distance is shorter than the actual distance. This bias is pronounced when measured through a conventional algorithm (e.g. compare panels a1 with a2; b1 with b2; c1 with c2 in Fig. 3) and when the radius is small (e.g. compare panels b1 with c1; b2 with c2 in Fig. 3). For *Simulation 3.1*, the novel algorithm was more reliable than the conventional algorithm as in *Simulations 2.1* and *2.2*.

In *Simulation 3.2*, the reliable range of speed estimation is relatively narrow compared to other simulations. Although the width of the reliable band is wider for the novel algorithm, the position of the band is different between the two algorithms. For example, for high values of the walking speed, the reliable band (dashed lines in Fig. 3e1, e2) lies along the time window of ca. 0.2 sec for conventional algorithm (Fig. 3e1) whereas it lies along ca. 0.3 sec for novel algorithm (Fig.3e2). There is no noticeable difference between conventional and novel algorithms for *Simulation 1*.

Throughout all the simulations, the geometric curvature is overestimated if speed is slow and the sampling window is short. This bias is present even when the actual geometric curvature is zero in *Simulation 1* where the actual trajectory is linear. As an exception, in Fig. 3c3 (upper right corner of the panel), there is an overestimation of geometric curvature for the specific combinations of straight speed and window length. This band seems to be induced when the travel distance between two observation points is the same as the perimeter of the circular path in the simulation, and hence it is particular to the circular path. Unlike the cases of the circular path, the estimated geometric curvature for sinusoidal-like paths is underestimated as the measured distance between two points gets longer (and time window gets longer).

The average standardized deviation of novel distance estimation was lower than conventional estimation given that speed estimation method is identical. For example, estimates from PtN_0.5_ exhibits lower standardized deviation compared to those from PtC_0.5_. Granted that conventional method of distance estimation was used, the fitting method of speed estimation is more accurate than the estimation from fixed time window. The overall standardized deviation was lowest when PtN_0.3_ was used for estimation, but the maximal ratio of standardized deviation (calculated as maximal standardized deviation divided by minimal standardized deviation among five simulations) was much greater for PtN_0.3_ (44, Table 2) compared to those values from FitC or FitN (5.95, 13.9, respectively, Table 2). This indicates that though the average standardized deviation *per se* might be smaller for PtN_0.3_, the stability of the estimation could be higher for FitC or FitN. Except for the PtN_0.3_, the standardized deviation of estimates from FitC or FitN are the lowest compared to other methods. The estimation of geometric curvature was most accurate if FitC or FitN methods were used but the difference of standardized deviation from FitC and Pt_0.3_ was not notable.

### 3.2. Estimation of empirical speed and geometric curvature using different methods

We applied the eight methods (FitC, FitN, PtC_0.1_, PtN_0.1_, PtC_0.3_, PtN_0.3_, PtC_0.5_, PtN_0.5_) to our empirical data composed of 104,508 trajectories from 39 video recordings filmed on an experimental arena in the field (Fig. 2d). We measured the mean and the standard deviation of corresponding speed and geometric curvature (Table 4). The differences among average estimates from FitC, FitN, PtC_0.1_, PtN_0.1_ were less than 2 mm/sec for speed and 0.01 mm^-1^ for geometric curvature. Though the sizes of the differences were minute, there was a significant difference in speed estimated from the four methods (Friedman test, *p* < 10^-10^). The same comparison was performed for geometric curvature. There was a significant difference among the geometric curvature measured by each method (Friedman test, *p* < 10^-10^). The average estimates of speed and geometric curvature decreased as window length increased. The mean adjusted *R*^2^ for FitC and FitN are 0.7843 and 0.7201, respectively, which differ significantly (Wilcoxon signed-rank test, *p* < 10^-10^). Additionally, it was revealed that the distribution of speed and geometric curvatures of the trajectories are inverted U-shaped (Fig. 5).

**Figure 5.**
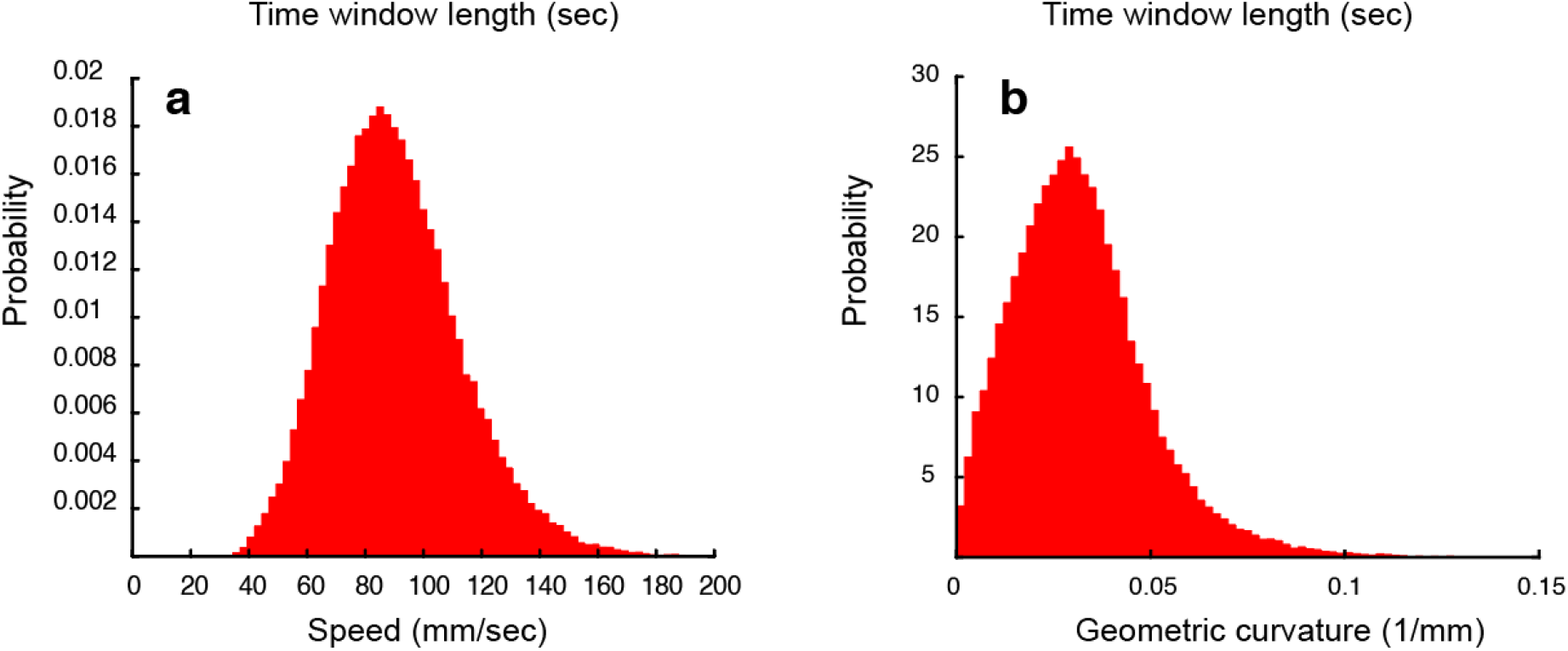
The distribution of speed and geometric curvature estimates from empirical data. **a, b**. The mean speed (a) and geometric curvature (b) obtained from 104,508 trajectories from 3 ant colonies using the FitN method follow unimodal distributions.

**Table 4.**
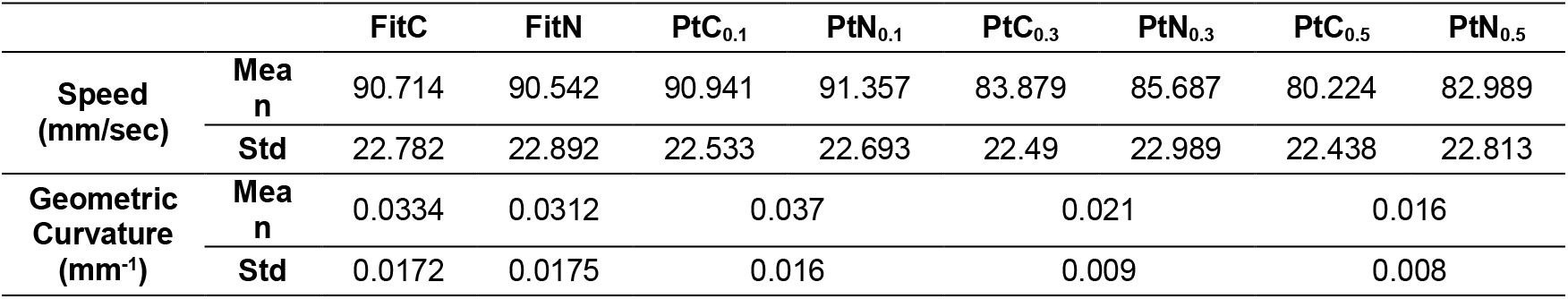
The mean and standard deviation of measured speed and geometric curvature of empirical data. Explanations of all methods are presented in Table 1.

|  |  | FitC | FitN | PtC<sub>0.1</sub> | PtN<sub>0.1</sub> | PtC<sub>0.3</sub> | PtN<sub>0.3</sub> | PtC<sub>0.5</sub> | PtN<sub>0.5</sub> |
| --- | --- | --- | --- | --- | --- | --- | --- | --- | --- |
| **Speed**<br>**(mm/sec)** | **Mean** | 90.714 | 90.542 | 90.941 | 91.357 | 83.879 | 85.687 | 80.224 | 82.989 |
|  | **Std** | 22.782 | 22.892 | 22.533 | 22.693 | 22.49 | 22.989 | 22.438 | 22.813 |
| **Geometric**<br>**Curvature**<br>**(mm<sup>-1</sup>)** | **Mean** | 0.0334 | 0.0312 | 0.037 |  | 0.021 |  | 0.016 |  |
|  | **Std** | 0.0172 | 0.0175 | 0.016 |  | 0.009 |  | 0.008 |  |

**Table 5.**
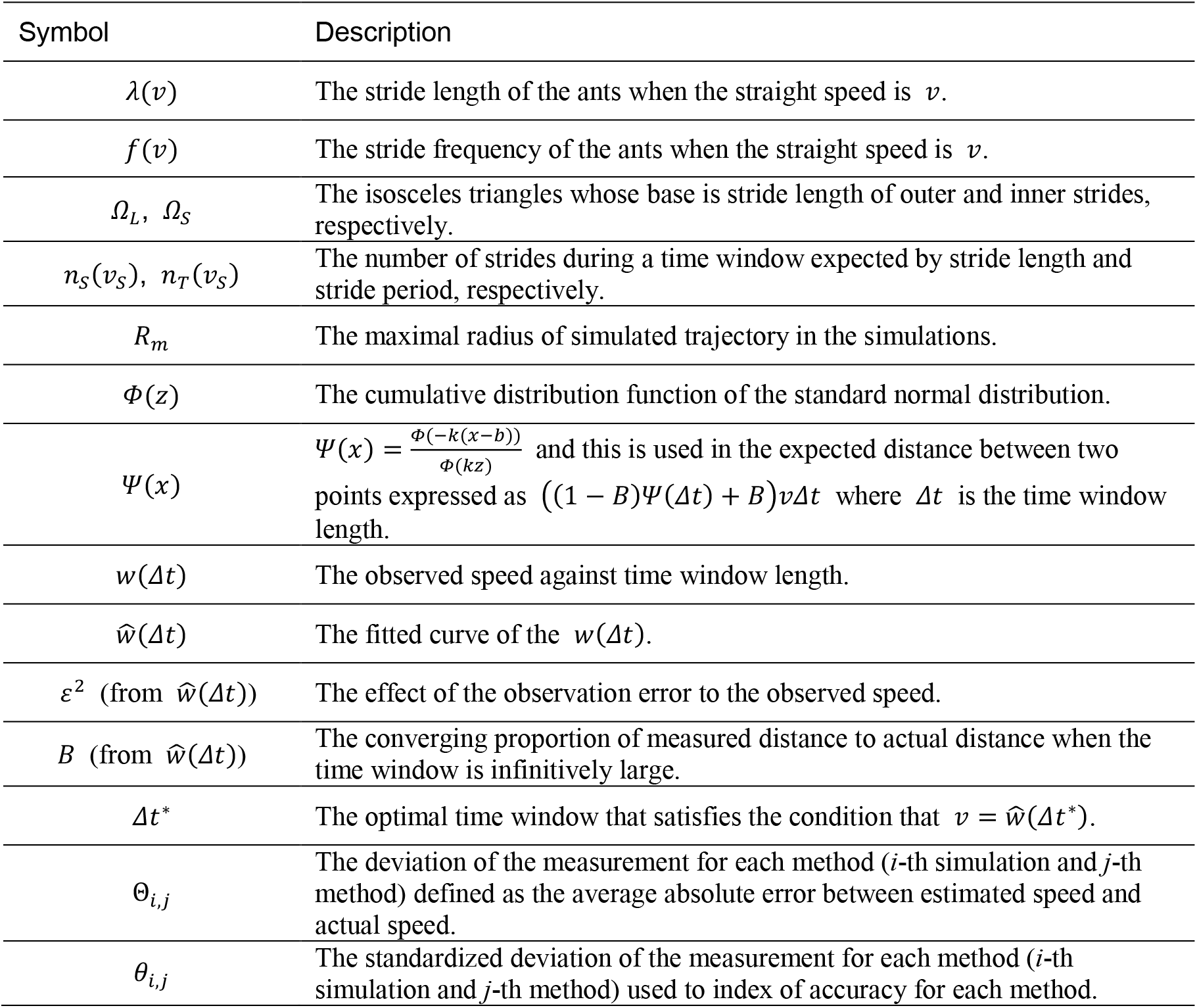
The list of variables and functions used in this study

## 4. Discussion

To overcome the complicatedness of the window length selection, many models have been developed^15–18^, but none of the models up to date has embraced the gait pattern of ants when estimating the travel distance. We propose in this study the algorithm and fitting method that can reliably estimate the speed and geometric curvature of ants’ trajectories by utilizing traits of ant gait.

The logical process in the model also provides the protocol for fitting speed with stride frequency and stride length. Presumably due to its simplicity, the stride frequency and length are often fitted linearly against the speed^13,14,21^ which yields a contradiction as shown in 2.1. Algorithm Explanation. Our novel equations for the relationships between stride length and speed and between stride period and speed remove this logical and mathematical contradiction, and therefore should be used in the future studies on ants. Those equation are also the basis for our novel method of speed estimations.

Unlike other stochastic algorithms for the time-invariant estimation of travel distance, our model utilizes the trend of estimated speed caused by change of time window length. The properties of the relationship between the time window (*x*-axis) and estimated speed (*y*-axis in Fig. 2c or Fig. 4c3-e3) may be used as indicators of measurement error and tortuosity. It allows us to estimate how error affect the results by inflating the measured distance and general tortuosity of the trajectory. For example, high values of measured speed when the time window is short compared to the other time windows indicate that the measurement error is relatively greater. Similarly, if the function representing the observed speed against time window (*w*(Δ*t*)) is relatively flat (i.e. The observed speed is almost invariant against window length as in Fig. 4c3) implies that the trajectory is straight, while a sharp decrease of observed speed as window length increases implies that the trajectory is highly convoluted (as in Fig. 4e3).

The decrease of observed speed with increasing time window is explained by the coastline paradox^23^. Though the actual path of ants would be curvilinear, due to the gap between frames, the recorded points are discretized, so that simple linking of the recorded points would yield a jagged path. This is supported by the fact that *ŵ*(*Δt*) is monotonically decreasing function, and average adjusted coefficient of determination (adjusted *R*^2^) of this regression to empirical trajectories is higher than 0.7. Due to the tradeoff of window length selection, researchers had to subjectively single out a specific window length in order to analyze and deduce the speed. For example Wang & Song (2016)^11^ selected 0.4 sec as the window length considering the fluctuation of the data points. Our algorithm enables researchers to automatically select the optimal time window length that is specific to each trajectory.

Theoretically, the geometric curvature should converge to zero for a directed walk as window length tends to infinity. As shown by the results of *Simulation 1*, though the actual line is perfectly straight, errors can falsely bring about the positive values of the curvature index.

Based on our results, we propose that in order to reliably estimate the speed of ants from empirical data researchers should measure coefficients of stride frequency 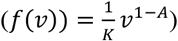 and stride length (*λ*(*v*) = *Kv^A^*) against walking speed by analyzing sample slow-motion videos. Applying the algorithm presented in this study would enhance the accuracy of the estimation as evidenced by simulation results, especially when the trajectories of ants are curved. The methods presented in this study would be useful to deduce speed and geometric curvature from videos. Other alternative algorithms with similar objectives presume low-frequency data collection from GPS (for example, wood turtles (*Glyptemys insculpta*) and white-nosed coati (*Nasua narica*), with maximal collection frequency of 4/min^15^), not suitable for data extracted from the videos of walking insect. As there are many indices of tortuosity suggested for animal behavior which are also affected by time window^5,24^, our algorithm can be utilized to select the most reliable tortuosity indices other than geometric curvature for ant trajectories, and other insects with similar gait characteristics (for example, common fruit fly (*Drosophila melanogaster*)^25,26^.

## Acknowledgments

Authors are grateful to Hyunwoo Goo for video processing and Hangah Lim for aid in field experiments.

## Appendix

Consecutive three points in a trajectory of an ant ((*X*(*t* – *δ*), *X*(*t*), *X*(*t* + *δ*)) = (*P*_1_, *P*_2_, *P*_3_)) were used to deduce a circumscribed circle, which was used as an osculating circle when calculating curvatures of trajectories. (These triplets are non-overlapping, and the points other than the triplets are not considered in the analysis.)

Suppose that the total angle between *P*_1_ and *P*_2_ (the first and second from the triplet) is *θ_T_* which is derived by 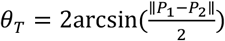. The number of strides (*n_S_*) is defined as the number of outer or inner stride during the time window. For example, if the time window length is the same as the period, then *n_S_* = 2 because an ant performs inner and outer strides during a period.

It is not certain which part at which a stride the body center *P*_1_ was measured. We can suppose either of four condition holds: (1) *P*_1_ is located at the middle of the inner stride; (2) *P*_1_ is located at the middle of the outer stride; (3) *P*_1_ is located at the end of the inner stride (equivalent, at the start of the outer stride); (4) *P*_1_ is located at the end of the outer stride (equivalent, at the start of the inner stride). We estimated the estimated speed from those four conditions and averaged the values to estimate the actual speed. If the speed of the body center is constant during the stride and the duration of inner and outer strides are the same, it can be expected that the probabilities that the actual body center is located at either of the aforementioned conditions are the same. Additionally, the total number of the strides would satisfy 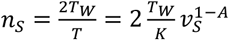.

### (1) Body center is at the middle of the inner stride

For a fixed *v_S_* and observed *R*, the outer stride length (equivalent to the stride length in a straight stride) is estimated as 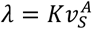. As assumed in the model, the height of the *Ω_S_* (the isosceles triangle with base *kλ*) is *L*.

Let the height of *T_L_* (the isosceles triangle with base *λ*) be *L*_0_. *L*_0_ should be short than *L* as *Ω_S_* and *Ω_L_* share the same hypotenuses but the base is longer for *Ω_L_*. (L: leg span)

**Figure A1.**
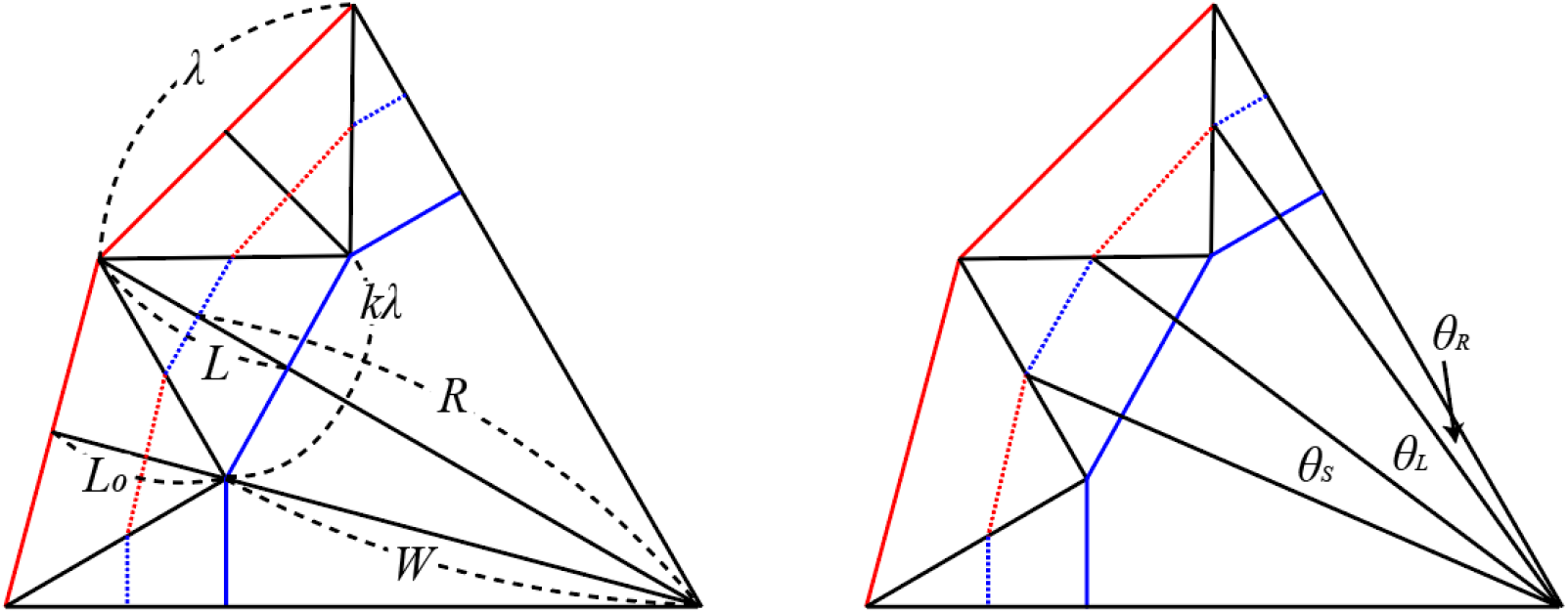

By conservation of triangle areas, Pythagorean theorem, and geometric symmetries, the followings hold:

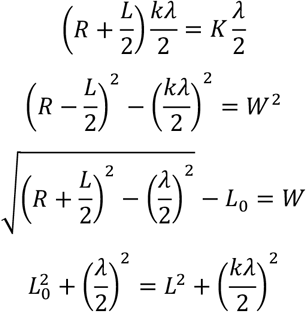

It is possible to computationally achieve *k, L*_0_, *K*, and *λ* that satisfy the aforementioned conditions. Then, the angle against the revolution center generated by the body center during the outer stride is 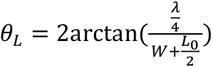 and that during the inner stride is 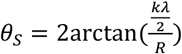. Define *θ_F_* = *θ_S_* + *θ_L_* which is the angle generated by an inner and outer stride. As it was assumed that *P*_1_ is located at the middle of the inner stride, three conditions are expected based on *θ_L_, θ_S_*, and *θ_T_*.

(1)-(i) *θ_T_* ≤ *θ_S_*/2 In this condition the total travel distance of the body center is 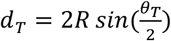, therefore the total number of the stride is 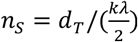.
(1)-(ii) 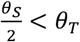 In this condition, the minimal number of the stride by first half inner stride is 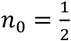. Define 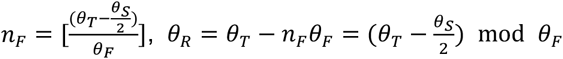 where [] is the floor function.

a. *θ_R_* ≤ *θ_L_* In this case, the length of body center movement after *n*_0_ + *n_F_* stride(s) is, by the law of cosines, 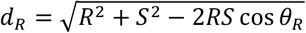 where *S* is the length of common hypotenuse shared by two isosceles triangles whose bases are *kλ* and *λ*, so that 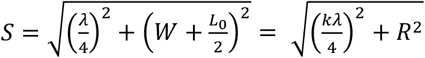. The addition of the number of progress by this movement is 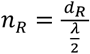 because the last stride during *P*_1_ and *P*_2_ is the outer stride. The total travel distance of the body center is 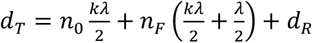.
b. *θ_R_* > *θ_L_* In this case, define *θ*_*R*2_ = *θ_R_* – *θ_L_*. Then, the length of body center movement after 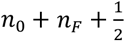 stride(s) is 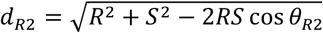. The addition of the number of progress by this movement is 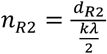 because the last stride during *P*_1_ and *P*_2_ is the inner stride. Define 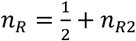. The total travel distance of the body center is 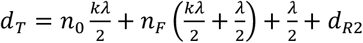. In both cases of (a) and (b), the total number of the stride *n_S_* = *n*_0_ + *n_F_* + *n_R_*. With given values of *θ_T_* and *R, n_S_* therefore can be represented as a function of *v_S_*. As the stride length is a differentiable and monotonically increasing function of *v_S_* as 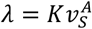, it is expected that *n_S_* by *ν_S_* is monotonically decreasing function of *ν_S_*. Therefore, *n_S_* can be derived spatially and temporal aspect. In a spatial aspect, *n_S_* is a function of *v_S_*. Let this relation be denoted by *F_S_*(*v_S_*) = *n*_0_ + *n_F_* + *n_R_*. (For brevity, assume that *n_F_* = *n_R_* = 0 and 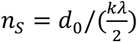 if *θ_T_* ≤ *θ_S_*/2.) In a temporal aspect, 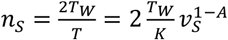. Let this relation be denoted by 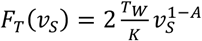. If *v_S_* is very small, then *λ* is infinitely small while and *T* is infinitely large, so that *F_S_*(*v_S_*) is infinitely large and *F_T_*(*v_S_*) is infinitely small, and *vice versa* when *v_S_* is very large. As *F_S_*(*v_S_*) is monotonically decreasing and *F_T_*(*v_S_*) is monotonically increasing, we can expect a single intersection point of *F_S_*(*v_S_*) and *F_T_*(*v_S_*) whose corresponding *v_S_* is the estimated straight speed. Based on this 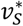, we can expect corresponding estimated values of *k, L*_0_, *K*, and *λ*. Again based on these values, we can expect the total estimated travel distance of the body center (*d_T_*) which would derive the actual speed of the body center on a curvy trajectory, *v_C_* = *d_T_*/*T_W_*. Let this value be denoted by *v*_*C*1_.

### (2) Body center is at the middle of the outer stride

With other notations being equal to case (1), by conservation of triangle areas, Pythagorean theorem, and geometric symmetries, the followings hold:

**Figure A2.**
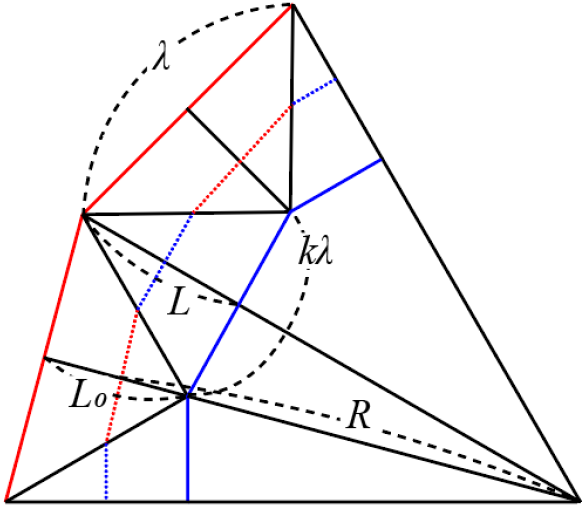

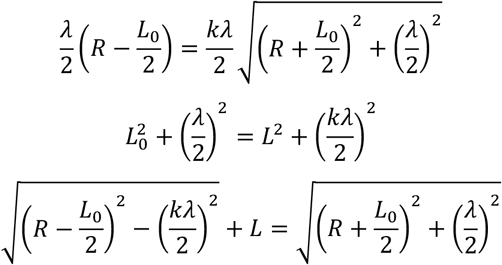

It is possible to computationally achieve *k, L*_0_, and *λ* that satisfy the aforementioned conditions. Then, the angle against the revolution center generated by the body center during the outer stride is 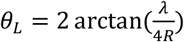 and that during the inner stride is 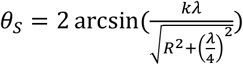. Define *θ_F_* = *θ_S_* + *θ_L_* which is the angle generated by an inner and outer stride. As it was assumed that *P*_1_ is located at the middle of the outer stride, three conditions are expected based on *θ_L_, θ_S_*, and *θ_T_*.

(2)-(i) *θ_T_* ≤ *θ_L_*/2 In this condition the total travel distance of the body center is 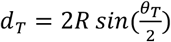, therefore the total number of the stride is 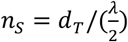.
(2)-(ii) 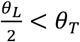 In this condition, the minimal number of the stride by first half inner stride is 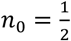. Define 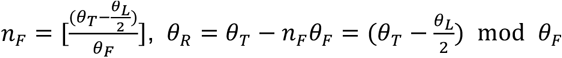.

a. *θ_R_* ≤ *θ_S_* In this case, the length of body center movement after *n*_0_ + *n_F_* stride(s) is, by the law of cosines, 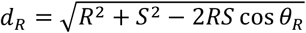 where *S* is the length of common hypotenuse shared by two isosceles triangles whose bases are *kλ* and *λ*, so that 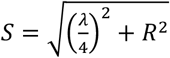. The addition of the number of progress by this movement is 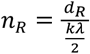 because the last stride during *P*_1_ and *P*_2_ is the outer stride. The total travel distance of the body center is 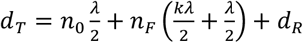.
b. *θ_R_* > *θ_S_* In this case, define *θ*_*R*2_ = *θ_R_* – *θ_S_*. Then, the length of body center movement after 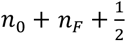 stride(s) is 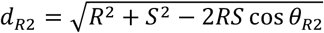. The addition of the number of progress by this movement is 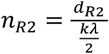 because the last stride during *P*_1_ and *P*_2_ is the outer stride. Define 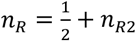. The total travel distance of the body center is 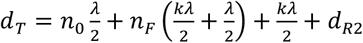. In both cases of (a) and (b), the total number of the stride *n_s_* = *n*_0_ + *n_F_* + *n_R_*. We can expect a single intersection point of *F_S_*(*v_S_*) and *F_T_*(*v_S_*) whose corresponding *v_S_* is the estimated straight speed. Based on this 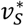, we can expect corresponding estimated values of *k, L*_0_, and *λ*. Again based on these values, we can expect the total estimated travel distance of the body center (*d_T_*) which would derive the actual speed of the body center on a curvy trajectory, *v_C_* = *d_T_*/*T_W_*. Let this value be denoted by *v*_*C*2_.

### (3) Body center is at the end of the inner stride

With other notations being equal to case (1), by conservation of triangle areas, Pythagorean theorem, and geometric symmetries, the followings hold:

**Figure A3.**
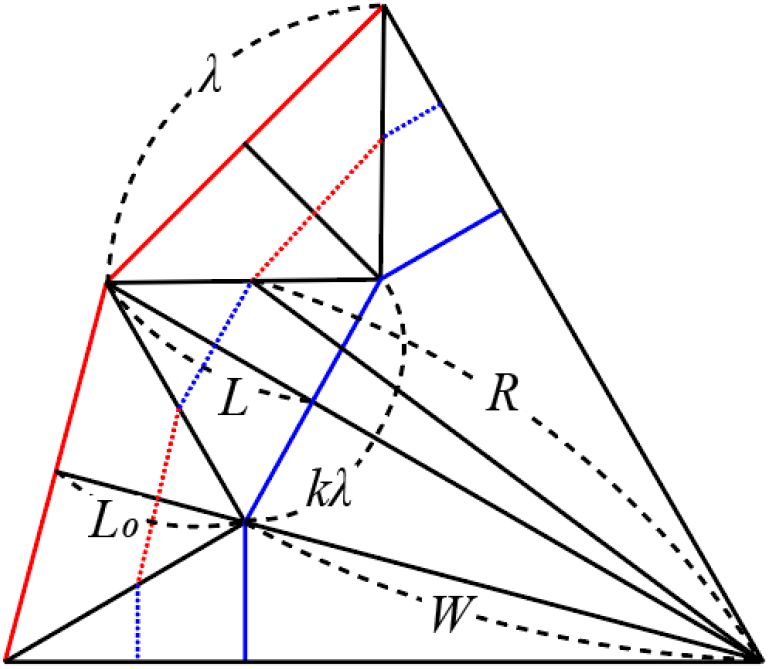

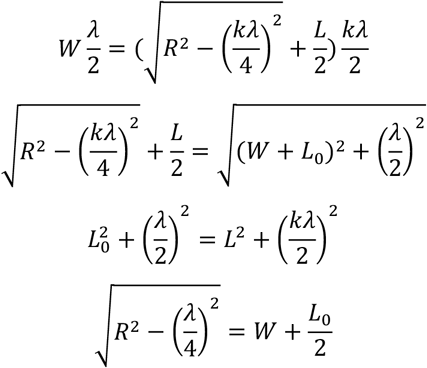

It is possible to computationally achieve *k, L*_0_, and *λ* that satisfy the aforementioned conditions. Then, the angle against the revolution center generated by the body center during the outer stride is 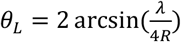 and that during the inner stride is 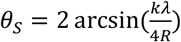. Define *θ_F_* = *θ_S_* + *θ_L_* which is the angle generated by an inner and outer stride. Define 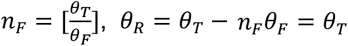 mod *θ_F_*.

As it was assumed that *P*_1_ is located at the middle of the inner stride, two conditions are expected based on *θ_R_, θ_L_, θ_s_*, and *θ_T_*.

1. *θ_r_* ≤ *θ_L_* Given this condition, 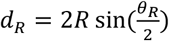 so that 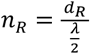 and *n_s_* = *n_F_* + *n_R_*. The total travel distance is 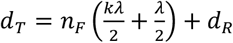.
2. *θ_R_* > *θ_L_* Given this condition, define *θ*_*R*2_ = *θ_R_* – *θ_L_*. 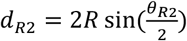 and 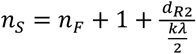. The total travel distance is 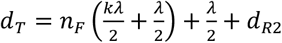. Based on these values, we can expect the total estimated travel distance of the body center (*d_T_*) which would derive the actual speed of the body center on a curvy trajectory, *v_C_* = *d_T_*/*T_W_*. Let this value be denoted by *v*_*C*3_.

### (4) Body center is at the end of the outer stride

All the values of *k, L*_0_, *λ, θ_L_, θ_S_* are the same as in (3). Define *θ_F_* = *θ_S_* + *θ_L_* which is the angle generated by an inner and outer stride. Define 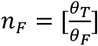, *θ_R_* = *θ_T_* – *n_F_θ_F_* = *θ_T_* mod *θ_F_*.

1. *θ_R_* ≤ *θ_S_* Given this condition, 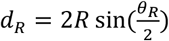 so that 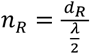 and *n_S_* = *n_F_* + *n_R_*. The total travel distance is 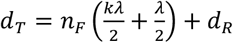.
2. *θ_R_* > *θ_S_* Given this condition, define *θ*_*R*2_ = *θ_R_* – *θ_S_*. 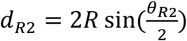 and 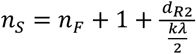. The total travel distance is 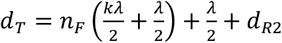.

Based on these values, we can expect the total estimated travel distance of the body center (*d_T_*) which would derive the actual speed of the body center on a curvy trajectory, *v_C_* = *d_T_/T_W_*. Let this value be denoted by *v*_*C*4_.

The estimated actual speed on a curved trajectory is 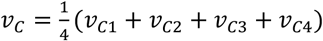.

